# CellFM: a large-scale foundation model pre-trained on transcriptomics of 100 million human cells

**DOI:** 10.1101/2024.06.04.597369

**Authors:** Yuansong Zeng, Jiancong Xie, Zhuoyi Wei, Yun Su, Ningyuan Shangguan, Shuangyu Yang, Chengyang Zhang, Wenbing Li, Jinbo Zhang, Nan Fang, Hongyu Zhang, Huiying Zhao, Yutong Lu, Jue Fan, Weijiang Yu, Yuedong Yang

## Abstract

The rapid evolution of single-cell sequencing technologies has facilitated precise transcriptomics profiling at the single-cell level, shedding light on the intricate heterogeneity within cellular populations. Despite these advances, the inherent diversity of cells and data challenges such as noise, batch effects, and sparsity, underscores the pressing need for a unified model to learn and represent cellular states effectively. Single-cell Large Language Models (LLMs) have been crafted to bridge this gap yet exhibit limited performance on human cells. This short-fall may stem from the confounding effects of training data from diverse species, partly because of limited cells for the single species. Here, we have compiled a dataset of approximately 100 million human cells sequenced by multiple technologies from human single-cell datasets with various file types deposited in public databases and websites. Leveraging these extensive data cohorts, we developed CellFM, a robust single-cell foundation model with an impressive 800 million parameters, marking an eight-fold increase over the current largest single-species model. To ensure the training of CellFM on the MindSpore AI framework from Huawei, we have integrated RetNet, a Transformer architecture variant with linear complexity for a balance between efficiency and performance, serving as the backbone of our model. Our comprehensive experiments have shown that CellFM outperforms existing models across diverse applications, such as cell annotation, perturbation prediction, and gene function prediction.

## 1 Introduction

Single-cell RNA sequencing (scRNA-seq) has ushered in a transformative era in molecular biology by facilitating the measurement of transcriptome profiles with an unprecedented scale and precision [1, 2]. With the progression of single-cell technologies, there has been a rapid accumulation of extensive single-cell datasets. However, the analysis of these datasets is fraught with significant challenges [3, 4], stemming from the inherent noise, sparsity, and batch effects present in the data. While numerous single-cell-specific tools [5–7] have been developed to tackle the analysis of these datasets, they often fall short in terms of performance when applied to new datasets and struggle to scale effectively with the growing size of data. More crucially, these tools do not fully exploit the rich information contained within the vast atlas datasets. Consequently, there is an imperative need to develop a unified model that can adeptly extract and encapsulate cellular states, capitalizing on the wealth of knowledge available in these abundant datasets.

To tackle this challenge, numerous single-cell Large Language Models (LLMs) have been developed to analyze single-cell data[8]. Benefiting from the impressive accomplishments of LLMs in natural language processing (NLP) [9, 10] and mitigating the training expenses, several researchers have explored the possibility of fine-tuning these models using a modest amount of single-cell data. For instance, Cell2Sentence [11] transforms the gene expression profiles of individual cells into sequences of gene names, arranged by their expression levels, and employs these sequences to fine-tune the GPT-2 model. Similarly, GenePT [12] and scELMo [13] harness GPT-3.5 to generate gene embeddings grounded in gene descriptions and metadata descriptors. While these methodologies have achieved enhanced performance through the fine-tuning of the GPT series, they do not yet fully exploit the rich gene expression data available in the expansive single-cell datasets.

To create single-cell LLMs from scratch, several gene-rank-based models are designed using the Transformer architecture [14]. For example, tGPT [15] captures the feature representation of transcriptomes through autoregressive modeling of a gene’s rank relative to its preceding neighbors. Specifically, tGPT takes a sequence of gene symbols ranked by their expression levels as input, aiming to predict the index of the subsequent gene in the dictionary based on the context of all previously mentioned genes. tGPT was trained on an extensive dataset comprising 22.3 million single-cell transcriptomes from humans and mice, showcasing enhanced performance across four distinct single-cell datasets. Likewise, Geneformer [16] employs a strategy that predicts gene positions within the cellular context to derive rank-based embeddings. Trained on an expansive dataset of 30 million single-cell transcriptomes encompassing a variety of human tissues, Geneformer has demonstrated exceptional predictive capabilities. However, the rank-based representation approach has its constraints. A significant drawback is that it does not fully utilize the gene expression values.

To address this gap, several bidirectional Transformer encoder-based single-cell Large Language Models (LLMs) have been crafted to fully use the detailed gene expression values. For example, scBert [17] divides the gene expression values of cells into multiple discrete “buckets,” effectively converting the continuous regression task of predicting gene expression into a discrete classification problem. Trained on millions of cells from humans, scBert has shown enhanced performance across various datasets. Similarly, scGPT [18] also segments gene expression but goes a step further by incorporating a specially designed attention mask mechanism to facilitate autoregressive prediction. Employing a self-supervised approach, scGPT simultaneously optimizes both cell and gene representations. Meticulously trained on a substantial collection of over 33 million cells from the human, scGPT has demonstrated impressive performance on a range of single-cell tasks. Diverging from the segmentation approach, scFoundation [19] directly forecasts raw gene expression values utilizing a mask autoencoder (MAE). scFoundation has been trained on an expansive dataset of about 50 million cells from humans with approximately 0.1 billion parameters, highlighting its ability to analyze single-cell data effectively. To fully use the prior information, GeneCompass [20] integrates four types of biological prior knowledge to deepen the understanding of gene regulatory mechanisms in a self-supervised manner. GeneCompass is pre-trained on a vast dataset consisting of about 50 million human cells and 50 million mouse cells, resulting in about 100 million parameters. It has proven its robustness and versatility across species, marking a significant advancement in single-cell analysis.

Though single-cell LLMs have been shown the potential to analyze single-cell data, they exhibit limited performance on human cells. This shortfall may stem from the confounding effects of training data sourced from diverse species. The scarcity of sufficient single-species training data, such as human cells, has constrained the development of single-species models with extensive parameters. The existing single-species models are typically limited to 50 million training data, resulting in model parameters of less than 100 million. One potential reason is that the single-cell datasets are difficult to collect. Since they are deposited everywhere with various types stored in formats such as FASTQ, matrix files, h5ad, Seurat objects, 10x Genomics formats, and compressed text matrices, to meet various data processing and analysis requirements.

To address this gap, we collect a lot of single-cell datasets from public databases and then make these data cleansing and standardization of unified formats, resulting in compiling a dataset of approximately 100 million human cells sequenced through various technologies. These datasets are twice as large as that of the current largest single-species model. Leveraging these extensive data cohorts, we introduce CellFM (Fig. 1), a robust single-cell foundation model with an impressive 800 million parameters, marking an eightfold increase over the current largest single-species model. To enhance the training of CellFM’s extensive parameters and to handle its substantial dataset, we have integrated ERetNet, a Transformer architecture variant with linear complexity. ERetNet’s design ensures a balance between efficiency and performance, serving as the backbone of our model. CellFM is developed using the MindSpore AI framework from Huawei and is trained on four Huawei Altas800 servers, each equipped with eight Ascend910 NPUs. Our comprehensive experiments have shown that CellFM outperforms existing models across diverse applications, such as cell annotation, perturbation prediction, and gene function prediction.

**Fig. 1.**
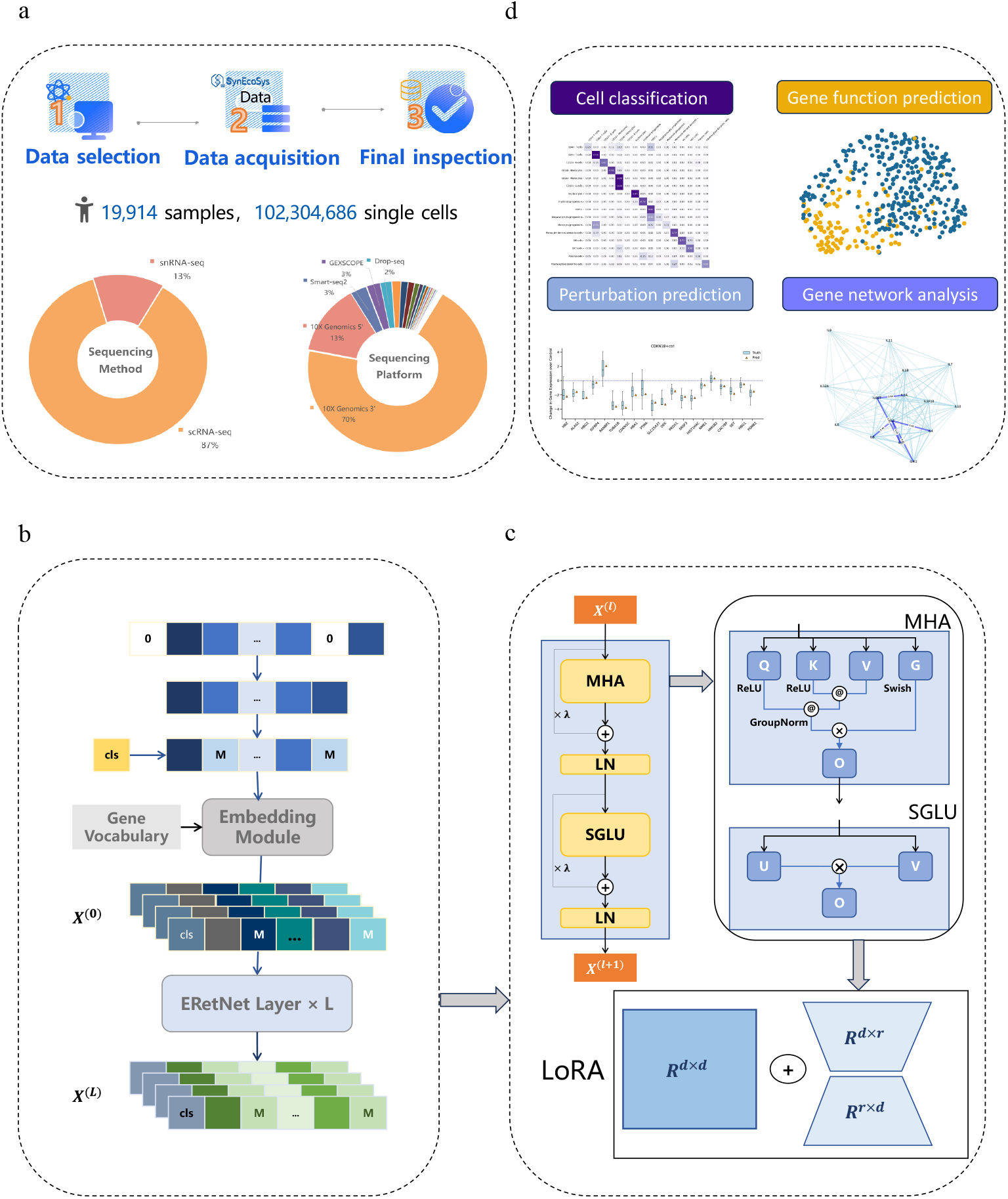
Overview of the CellFM Framework. (a) The collecting workflow and constituents of the training dataset employed within CellFM. (b) CellFM consists of the Embedding module, the ERetNet module, and the LoRA module. The expression of the cell is first fed into the Embedding module to obtain the initial token values of each gene. The embedded gene tokes of the cell are then inputted into the ERetNet module to learn the gene-to-gene relationships and gene embeddings. Finally, the Low-Rank Adaptation (LoRA) is implemented to minimize the number of training parameters of CellFM. (c) Each ERetNet Layer block integrates Multi-Head Attention (MHA), SGLU activation, and Layer Normalization (LN). (d) The pre-trained CellFM model is adaptable for a multitude of single-cell downstream analyses including cell type annotation, perturbation prediction, gene network inference, and gene function prediction.

## 2 Results

### 2.1 Overview of CellFM

Single-cell sequencing technology is crucial for revealing the detailed landscape of cellular diversity and function at the individual cell level. As single-cell sequencing technologies continue to evolve, a vast array of datasets has been amassed, laying a robust groundwork for the training of single-cell LLMs. However, these datasets are available on numerous public platforms including NCBI GEO [21], ENA [22], GSA [23, 24], and ImmPort [25], we thus have meticulously curated original single-cell data from these esteemed public databases (Fig. 1a). These datasets are stored in multiple formats, such as Fastq data, expression matrices, or Seurat/Scanpy objects. We process raw fastq data into the gene expression matrix through primary analysis software provided by manufacturers. Subsequently, all acquired expression matrices were processed using a standardized data analysis workflow facilitated by the SynEcoSys® single-cell database (Singleron Biotechnologies) [26]. This process involved three key steps including quality control for filtering cells and genes, gene name standardization according to HUGO Gene Nomenclature Committee (HGNC) guidelines, and converting the data to a unified sparse matrix format for subsequent analysis. Our efforts have successfully aggregated 19,914 samples, totaling 102,304,686 human cells sequenced through various single-cell sequencing technologies such as 10x, Smart-seq2, and Drop-seq.

Building on our comprehensive collection of human cell data, we introduce CellFM, an efficient foundation model endowed with 800 million parameters, designed to streamline the analysis of single-cell data (Fig. 1b). The model’s core is comprised of a series of stacked ERetNet Layers, each equipped with multi-head attention mechanisms that concurrently learn gene embeddings and the complex interplay between genes. CellFM begins by converting scalar gene expression data into rich, high-dimensional embedding features through its embedding module. These gene embeddings are then fed into L ERetNet Layers, which are adept at capturing the nuanced relationships among genes based on their expression profiles. Each ERetNet Layer is composed of several key components: the Gated Multi-head Attention (MHA) unit, the Simple Gated Linear Unit (SGLU), and Layer Normalization (LN). Collectively, these elements empower the ERetNet Layer to achieve training parallelism, cost-effective inference, and superior performance (Fig. 1c). Furthermore, CellFM integrates a low-rank adaptive module (LoRA), which serves to reduce the number of trainable parameters during the fine-tuning phase when adapting the model to new datasets. Once pre-trained, CellFM is versatile and can be applied to a multitude of single-cell downstream applications, such as gene function prediction, cell type annotation, perturbation effect prediction, and gene network analysis (Fig. 1d).

### 2.2 CellFM improves the accuracy of gene function prediction

Gene function prediction is a cornerstone for deciphering the roles and properties of genes under diverse conditions [16]. With the human genome comprising approximately 20,000 protein-encoding genes [27], and a significant portion lacking functional annotations, the accurate prediction of gene functions is imperative for a deeper understanding of their roles within biological systems. In this study, we focused on predicting three distinct gene function categories including Dosage sensitivity (referred to as T1), Bivalent methylation status versus non-methylated (T2), and Bivalent methylation versus Lys4-only methylation (T3). These categorizations represent binary classification challenges, where model predictions are assessed against actual gene function labels. Since the limited number of genes in these three tasks, typically fewer than 1000, fine-tuning existing models presents a challenge. To make a fair comparison, we adopted a zero-shot learning strategy for each model on the gene function prediction task. As shown in Fig. 2a, our model demonstrated remarkable results, achieving an average accuracy score (ACC) of 86.1% on the T1 task. This performance surpasses existing methods, with a notable 34% and 37.1% increase over scGPT and Geneformer, respectively. This trend of enhanced performance was consistent across the T2 and T3 tasks. Further analysis through macroF1 values revealed that our approach out-performed its competitors, with improvements of 31.2% and 43.9% over scGPT and Geneformer, respectively (Fig. 2b). While scGPT and Geneformer showed comparable ACC values across the tasks, scGPT excelled over Geneformer in macro F1 scores. The superior performance of our model was substantiated by visualizations using Uniform Manifold Approximation and Projection (UMAP) on the gene embeddings obtained from each pre-trained model. Our model’s ability to distinctly categorize dosage-sensitive from non-sensitive genes was evident in Fig. 2c, unlike scGPT and Geneformer, which exhibited an overlap in the embedding space that could undermine gene function prediction accuracy. Similar distinctions were observed in the T2 and T3 tasks. In summary, our findings underscore the model’s proficiency in accurately predicting gene functions utilizing a zero-shot approach, showcasing its efficacy without the need for extensive model fine-tuning.

**Fig. 2.**
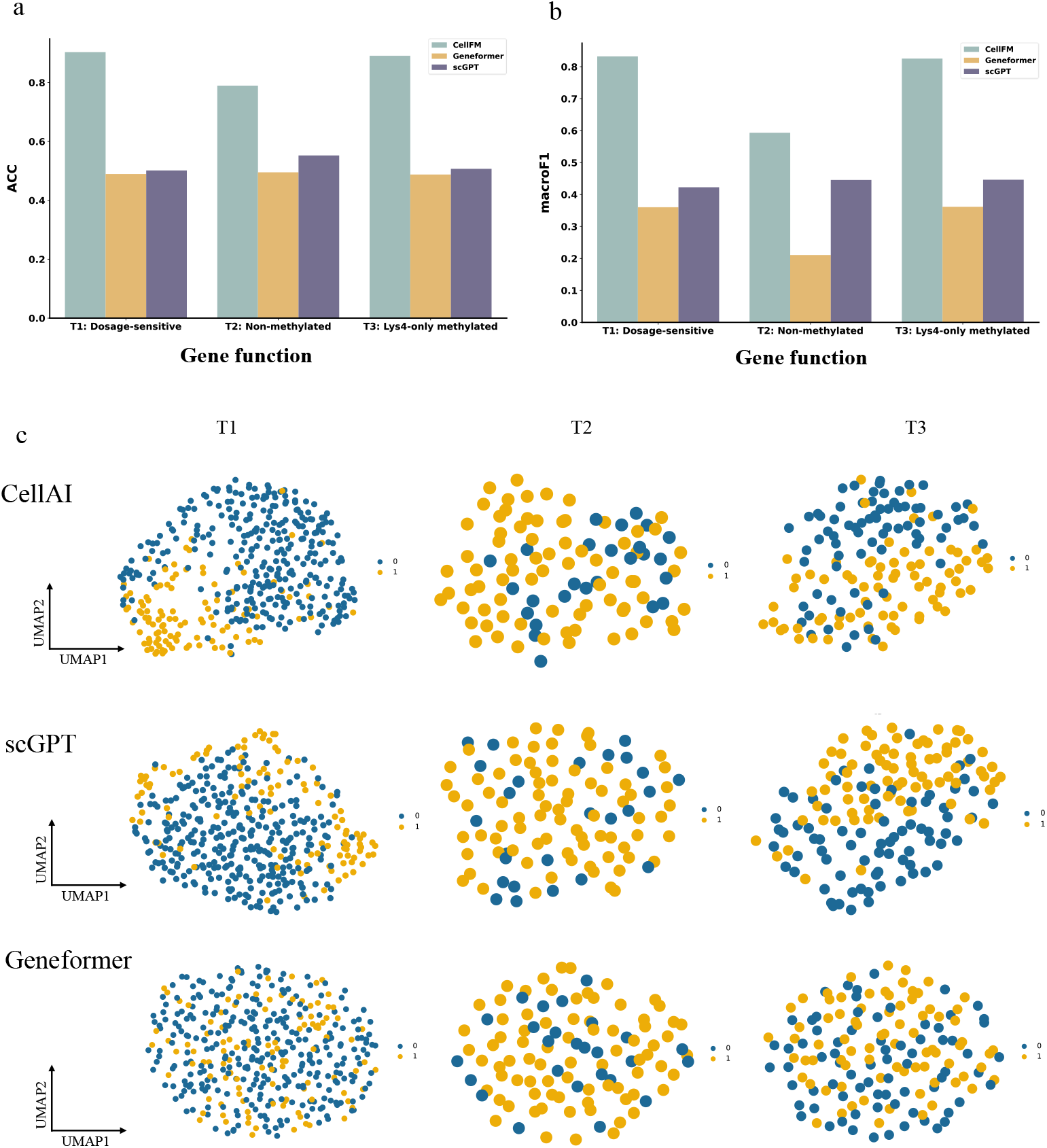
Comparison of Gene Function Prediction performance in a zero-shot setting: (a) accuracy (ACC) scores and (b) Macro-F1 values for CellFM, Geneformer, and scGPT. (c) The visualization results of each model are plotted by the UMAP using the gene embedding generated through each model.

### 2.3 The performance of CellFM on perturbation response and reverse perturbation prediction

Recent sequencing and gene editing breakthroughs have opened doors to large-scale perturbation experiments, allowing for detailed characterization of how cells respond to various genetic alterations. Yet, the vast array of potential genetic modifications poses a challenge that lies beyond the scope of feasible experimentation. To transcend this hurdle, CellFM has been deployed to harness insights from cellular reactions observed in known experiments and project them to foresee responses to previously unexplored perturbations. Similar to the gene function prediction task, we used the zero-shot strategy for perturbation prediction. Initially, we harnessed CellFM to generate cell embeddings for every perturbation dataset. These embeddings were subsequently utilized to develop a Multi-Layer Perceptron (MLP) model. Specifically, the MLP model ingests the embeddings from control cells that have not been altered and outputs the embeddings of the cells post-modification. The optimization of the MLP module is guided by minimizing the Mean Squared Error (MSE) between the predicted state of the modified cell and the actual state of the perturbed cell. Consequently, once trained, the MLP model is adept at forecasting the cellular responses to modifications by relying on expression data from control genes.

To systematically assess CellFM’s proficiency in predicting perturbations, we utilized two Perturb-seq datasets in our study: (1) the Adamson dataset, encompassing 87 single-gene perturbations with roughly 100 cells per perturbation and at least 7,000 control cells; and (2) the Norman dataset, which includes 131 dual-gene and 105 single-gene perturbations. As depicted in Fig. 3a, we employed the Pearson correlation metric on the top 20 differentially expressed genes (De) to evaluate each model, where Δ denotes the degree of gene expression alteration post-perturbation relative to the control state. CellFM excelled in this evaluation, achieving results that surpassed the second-ranked method scGPT by 1.65% on the Adamson dataset and 2% on the Norman dataset in terms of the average values, respectively. Additionally, we assessed the models’ performance by calculating the mean square error (MSE) for the top 20 differentially expressed (De) genes, comparing predicted post-gene expression profiles to the actual ones. As shown in Fig. 3b, CellFM consistently achieved lower MSE values across all perturbation datasets, outperforming the scGPT and MLP baseline models. To substantiate CellFM’s superior performance, we presented visual predictions for two specific perturbation cases from the Adamson dataset in Fig. 3c. Here, CellFM demonstrated its ability to accurately forecast the expression trend changes for all top 20 DE genes.

**Fig. 3.**
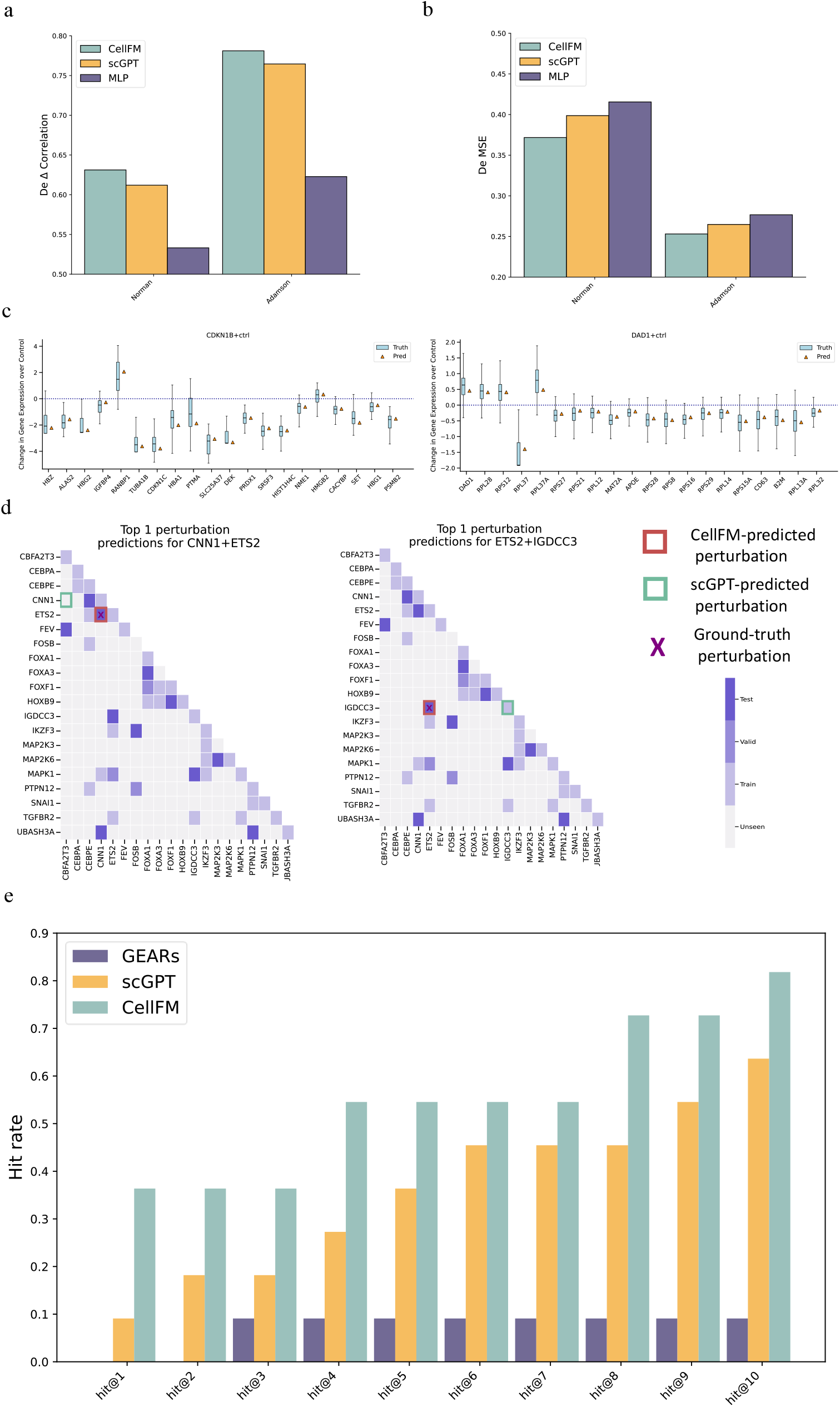
Analysis of Perturbation Response and Reverse Perturbation Predictions. (a) In a zero-shot learning context, a comparative analysis of CellFM against other models on the Norman and Adamson datasets, emphasizing the Pearson correlation coefficient between the predicted and actual changes in gene expression for the top differentially expressed genes. (b) The mean square error between predicted and actual post-gene expressions for the top 20 differentially expressed (DE) genes. (c) Examination of two specific perturbations involving genes CDKN1B and DAD1. The visualization displays the predicted versus actual gene expression changes for the top 20 DE genes. The box plots indicate the interquartile range of expression change, with the median represented by the central line. Whiskers extend to 1.5 times the interquartile range, and horizontal dashed lines indicate the null effect on gene expression. (d) A graphical representation of potential perturbation combinations across a 20-gene space, differentiated by experiment type (train, valid, test, unseen). Predicted perturbations are indicated by square boxes, with the actual source perturbation marked by a cross. (e) The accuracy of each model in predicting the correct source of perturbation among the top 10 predictions for eleven test cases.

Beyond forecasting the outcomes of gene perturbations, the accurate prediction of CRISPR target genes that prompt cellular recovery from disease states is equally significant. This challenge, termed in silico reverse perturbation prediction by scGPT, entails determining the original genetic perturbation from a given resultant cell state. To substantiate the versatility of our model for reverse perturbation prediction, we executed in silico experiments utilizing the Norman dataset. Specifically, we concentrated on a subset of the Norman dataset that included perturbations of 20 key genes. This subset encompasses a combinatorial space of 210 potential one-gene or two-gene perturbation combinations. We fine-tuned CellFM on 18% of these known perturbations and subsequently evaluated the model’s ability to predict the source of perturbations for unseen altered cell states. CellFM demonstrated remarkable success, accurately predicting the perturbations that would yield the observed cellular outcomes. For instance, it correctly identified the combination of CNN1 and ETS2 genes as the top prediction for one test case (illustrated in Fig. 3e). Comparable accuracy was observed for the perturbation involving ETS2 and IGDCC3 genes. In aggregate, CellFM achieved an average of correctly identified perturbations in 81.8% of the top 10 predictions (Fig. 3f). The performance of our model notably exceeds that of the scGPT method, which is ranked second, by a substantial improvement of 18.1%. As the number of top predictions considered increased, both CellFM and scGPT demonstrated enhanced performance. However, GEARs did not exhibit a comparable level of improvement in accuracy. These findings underscore the superior performance capabilities of single-cell LLMs compared to the model with small parameters.

### 2.4 Cell type annotation with CellFM

Cell identification is a cornerstone of single-cell data analysis, essential for uncovering the cellular heterogeneity within biological samples. To evaluate CellFM’s competency in cell type annotation, we conducted an exhaustive benchmark against several leading models. This assessment spanned both intra- and cross-dataset evaluations, employing a zero-shot learning approach. Our initial evaluation involved eight intra-datasets, each comprising a single batch of data. Following the methodology established by scGPT [18], we segmented each intra-dataset, allocating 70% for training and the remainder for testing. As illustrated in Fig. 4a, CellFM excelled baselines in terms of ACC across intra-datasets. The average ACC for CellFM was 68.8%, surpassing the second-ranked model Geneformer by 10.8%. A similar trend could be found when measured by the MacroF1 values (Fig. 4b). To substantiate the superior outcomes of CellFM, we present a case study of the immune dataset. The predictions are visualized in Fig. 4c-d. CellFM achieved high precision for most cell types except for a few rare cell types including Plasma cells and Monocyte progenitors. In contrast, scGPT only achieved accurate prediction on the major cell types including CD4T+ cells and CD14+ Monocytes.

**Fig. 4.**
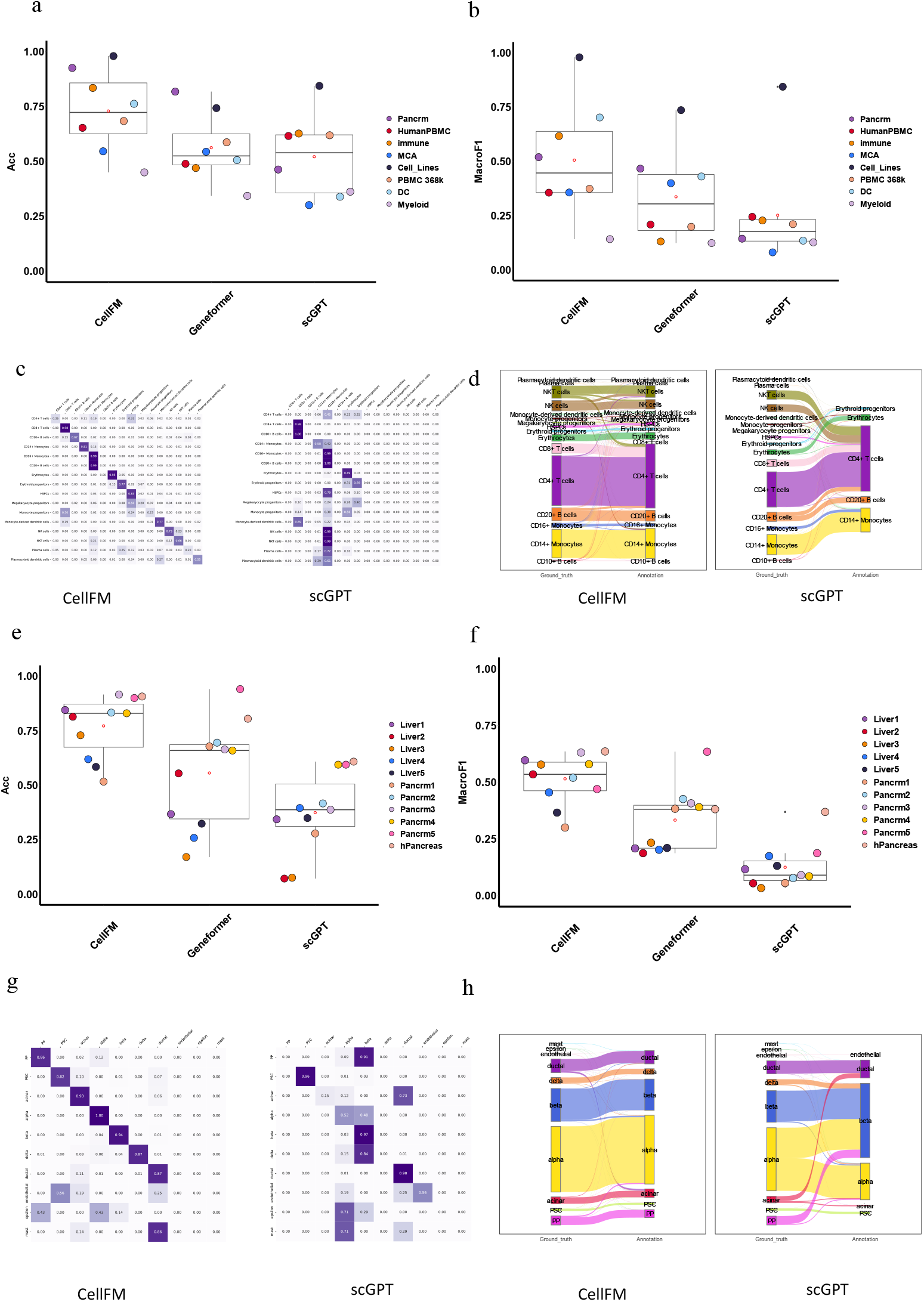
Zero-shot cell type annotation performance of each model. (a) Boxplots summarize the ACC scores for each method on n=8 biologically independent paired intra-datasets. (b) Boxplots summarize the Macro-F1 scores for each method on n=8 biologically independent paired intra-datasets. (c) The confusion matrix between predicted and annotated cell types in the immune dataset. (d) River plots illustrate the predicted cell types and their relationships to the actual cell types on the immune dataset. (e) Boxplots summarize the ACC scores for each method on n=11 biologically independent paired inter-datasets. (f) Boxplots summarize the Macro-F1 scores for each method on n=11 biologically independent paired inter-datasets. (g) The confusion matrix between predicted and annotated cell types in the hPancreas dataset. (h) River plots illustrate the predicted cell types and their relationships to the actual cell types on the hPancreas dataset.

To substantiate the efficacy of our model in the context of batch effects, we assessed its performance on 11 paired cross-batch datasets. In each scenario, a distinct batch of data was designated as the test set, with the remaining data constituting the training set. As depicted in Fig. 4e-f, our model continued to outperform its competitors. Specifically, it achieved an average ACC and Macro F1 that were 21.6% and 18.2% higher, respectively, than those of the second-ranked method, Geneformer. Notably, all rival models struggled with the Liver4 batch from the Liver dataset, recording an accuracy below 40%. In contrast, our model showcased its resilience and robustness by attaining a remarkable 62% accuracy. To further attest to the exceptional performance of our model, we present an illustrative case study of the hPancreas dataset. The predictions are graphically represented in Fig. 4g-h. Impressively, CellFM achieved a high precision of over 0.8 for the majority of cell types. The sole exceptions pertained to the rare cell types that were sparsely represented within the test subset, including mast cells, epsilon cells, and endothelial cells. Each of these cell types constituted less than 5% of the total cell population.

### 2.5 Deciphering gene relationships with CellFM

The intricate interplay among target genes within a Gene Regulatory Network (GRN) is pivotal for orchestrating key biological processes. Here, we aimed to substantiate the biological relevance of the gene networks delineated by CellFM and assess their utility in uncovering gene programs. To this end, we evaluated CellFM’s proficiency in identifying GRNs by clustering functionally related genes using the K-nearest neighbors algorithm based on gene token embeddings. As illustrated in Fig. 5a, CellFM successfully identified a cohort of genes with substantial similarity scores, such as IL2, IL4, and IL6. These genes, as it turns out, are integral to the Th17 cell differentiation pathway [28]. Specifically, IL-2, IL-4, and IL-6 exert significant influence over the Th17 cell differentiation process, thereby playing a crucial role in immune response modulation and the pathogenesis of autoimmune diseases. In summary, our findings demonstrate that CellFM is adept at identifying functionally coherent gene groups, thereby highlighting its potential as a valuable tool for gene network analysis and discovering novel gene programs.

**Fig. 5.**
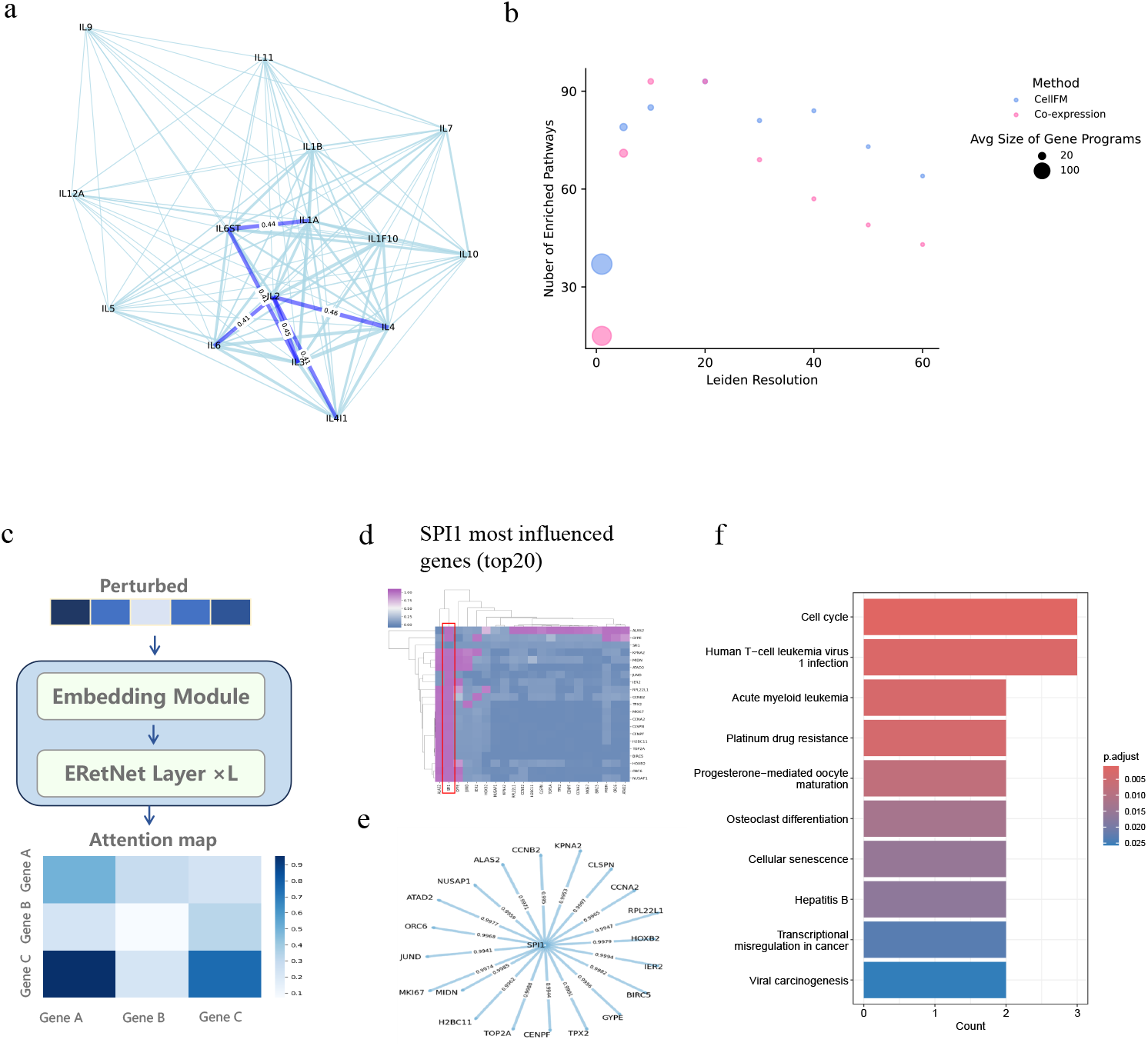
Gene-Gene Relationships Unveiled by CellFM. (a) A gene cluster comprising IL2, IL4, and IL6 was identified through the cosine similarity of gene embeddings generated by the zero-shot CellFM. (b) This analysis compares the number of enriched pathways derived from gene programs extracted by CellFM and a coexpression network within an immune-related human dataset, across various Leiden clustering resolutions. (c) This workflow outlines the process of identifying the most influenced genes through attention maps, where attention scores from perturbed cell states are sequentially ranked to select the most impacted genes. (d) The heatmap displays the connectivity changes in the network of the top 20 genes most influenced by the repression of SPI1. (e) The network graph represents the top 20 genes, with ChIP-seq predicted targets validated in the ChIP-Atlas database, highlighted in light blue. (f) The heatmap illustrates the pathways mapped using the Kyoto Encyclopedia of Genes and Genomes (KEGG) database, based on the top 20 genes most significantly impacted by SPI1.

To rigorously verify the identified gene programs, we conducted a comprehensive pathway enrichment analysis using the Reactome database (https://reactome.org/). Concretely, we meticulously selected and categorized gene programs based on the gene embeddings generated by CellFM. This process was facilitated by applying the Leiden clustering algorithm across different resolutions. As illustrated in Fig. 5b, we juxtaposed the results yielded by CellFM with those obtained from co-expression network analysis. CellFM consistently revealed a significantly higher number of enriched pathways across all clustering resolutions, except for the resolution at 10. To further validate the efficiency of the pathways identified by CellFM, we conducted a comparative analysis of the pathways between CellFM and the co-expression network at a resolution of 40. Both methodologies identified 25 common pathways. CellFM uniquely identified an additional 59 pathways, 7 of which were pertinent to immune system processes. Conversely, the co-expression network uniquely identified 32 pathways, of which only 2 were associated with immune functions. These comprehensive findings underscore CellFM’s superior capacity to capture subtle and intricate gene-gene interactions, thereby enabling the elucidation of specific biological mechanisms within a broader cellular context.

To further substantiate our model can also identify gene relationships involved in perturbation experiments, CellFM discerns the genes most markedly affected by the perturbations by scrutinizing the attention maps (Fig. 5c). This approach affords a deeper understanding of context-specific gene regulatory interactions within individual cells. For instance, we examined the perturbed gene SPI1 within the Adamson dataset as a case study. As illustrated in Fig. 5d, CellFM identified the top 20 genes most influenced by the repression of SPI1, a gene encoding a transcription factor. These genes were corroborated as signaling targets of SPI1, as evidenced in the ChIP-Atlas database [29] (Fig. 5e). Furthermore, CellFM was able to capture distinct pathway-activation patterns (Fig. 5f) through the genes most influenced by SPI1 [30–32]. These attention-based insights further substantiate the gene network learned by CellFM at the cellular level, augmenting the interpretability of the biological mechanisms captured by the model.

## 3 Discussion

To aid efficient analysis of the single-cell data and harness the wealth of knowledge contained within single-cell atlas datasets, we have introduced a state-of-the-art foundation model known as CellFM. This model was pre-trained on our meticulously curated datasets, encompassing about 100 million human cells. These datasets empower CellFM to generate an expansive set of 800 million model parameters, marking an eightfold increase over the parameters present in the current single-cell models trained on a single species. To augment the training efficiency of CellFM, we have adopted the ERetNet architecture as its core. This network represents an advancement over the traditional RetNet framework, offering enhanced parallel training capabilities and cost-effective inference. These features collectively contribute to CellFM’s exceptional performance. Moreover, CellFM incorporates a Low-Rank Adaptive module designed to minimize parameter count during fine-tuning, thereby optimizing the model for specific tasks without compromising its generalizability. Through a series of comprehensive experiments, CellFM has demonstrated its effectiveness across a range of single-cell tasks including cell type annotation, prediction of responses to perturbations, gene network analysis, and gene function prediction.

Previous single-cell Large Language Models (LLMs) have demonstrated promising capabilities across various tasks. However, most of these models were trained on data encompassing multiple species. These settings limited the performance of these models on a single species such as humans since the confounding effects of training data sourced from diverse species. The scarcity of sufficient single-species training data has constrained the development of LLMs with extensive parameters. The existing single-species models are typically limited to 50 million training data, resulting in model parameters of less than 100 million. To address this gap, we have compiled a dataset comprising about 100 million human cells, specifically to refine our model’s training. This expansive dataset empowers CellFM to effectively learn rich prior knowledge, thereby enabling the creation of its advanced model with 800 million parameters. CellFM’s remarkable zero-shot performance in gene function prediction tasks reinforces the effectiveness of our model.

To satisfy the model’s training on large-scale datasets, we have chosen a variant of the RetNet architecture as CellFM’s foundation, diverging from the Transformers used in other single-cell LLMs. The RetNet architecture facilitates parallel, recurrent, and chunkwise processing, which we have refined by integrating the SGLU module, amplifying training efficiency. Additionally, we have embedded the Low-Rank Adaptive (LoRA) strategy within CellFM, optimizing its training on new datasets with similar characteristics. The combination of this efficient training architecture and the comprehensive datasets forms the basis for developing the current largest CellFM model, equipped with 800 million parameters. CellFM is developed using the MindSpore AI framework from Huawei and is trained on four Huawei Altas800 servers, each equipped with eight Ascend910 NPUs. Our rigorous experiments demonstrated the model’s adaptability and potency in multiple single-cell downstream tasks. In the spirit of research collaboration, we are dedicated to sharing our progress by making the CellFM codes and the pre-trained model publicly available. This initiative aims to provide researchers with a unified framework that streamlines the adoption of pre-trained models for their distinct research goals.

To enhance the capabilities of our model, we have outlined several optimization strategies for CellFM. Initially, we aim to expand the diversity and scope of the training data by integrating multi-omic datasets, spatial data, and samples representing various disease states. This enrichment will facilitate a more comprehensive grasp of the underlying biological mechanisms. Additionally, leveraging prior knowledge of gene relationships, such as those involved in signaling and functional pathways, has demonstrated the potential to bolster the performance of single-cell data analysis [33]. Consequently, we plan to explore the integration of these gene interactions in the forthcoming stages of CellFM’s development. Lastly, in the interest of advancing the interpretability of our machine learning model, we are committed to incorporating explainable AI methods. By doing so, we aim to provide users with insights into the decision-making process of the model. This approach not only promotes transparency but also fosters greater confidence in the reliability and accuracy of CellFM’s predictions.

## 4 Methods

### 4.1 Data collection

All training data utilized in this study were sourced from reputable public databases. Specifically, from April 2021 to August 2023, we identified datasets leveraging keywords like “single-cell RNA sequencing,” “single-cell transcriptome,” and “single-cell RNA.” These keywords were used to search through databases such as NCBI GEO [21], ENA [22], GSA [23, 24], ImmPort [25], and others. In our selection process, we carefully curated the datasets, retaining only those human single-cell datasets that were relevant to our study. These datasets were encountered in multiple formats, including Fastq data, expression matrices, and Seurat/Scanpy objects. Our initial step involved transforming the raw Fastq data into expression matrices using primary analysis software supplied by the manufacturers. Following this, all obtained and transformed expression matrices underwent pre-processing through a standardized workflow provided by the SynEcoSys® single-cell database from Singleron Biotechnologies [26]. This workflow included several critical steps: (1) Quality control involved filtering cells based on a minimum gene count threshold of 200 genes per cell; (2) Gene name standardization was conducted in accordance with the HUGO Gene Nomenclature Committee (HGNC) guidelines, ensuring that gene aliases in each dataset were converted to their respective HGNC-approved gene symbols. This step guaranteed the uniqueness and consistency of gene names across all datasets. Finally, the expression matrices for each sample were converted into a unified sparse matrix format, preparing them for subsequent model training.

### 4.2 CellFM architecture

The CellFM model comprises three core components including the embedding module, the ERetNet module, and the LoRA module (Fig. 1). The embedding module in CellFM maps one-dimensional scalar values of gene expression to high-dimensional embedding features for model training, enabling the representation of gene expressions in a high-dimensional space. CellFM then applies the ERetNet module to learn the relationships among genes based on the gene expression information. In parallel, CellFM uses the LoRA module to help train CellFM more efficiently by reducing the number of parameters when adjusting model weights with new data.

### 4.3 The embedding module

To efficiently train CellFM, we have set an upper limit on the number of genes it inputted, defined by the threshold *l*_*max*_ = 4096. For each cell, if the number of expressed genes exceeds *l*_*max*_, we randomly select *l*_*max*_ genes with high expression values. Conversely, if a cell has fewer expressed genes than *l*_*max*_, we pad the gene IDs and fill them with random values to extend the gene length to *l*_*max*_. We then apply a Multilayer Perceptron (MLP) to map the scalar expression values of genes to embedding vectors necessary for the ERetNet module as follows:

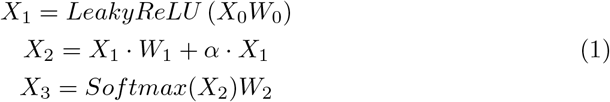

where *W*_0_ ∈ℝ^1×*b*^, *W*_1_ ∈ℝ^*b*×*b*^, and *W*_2_ ∈ℝ^*b*×*d*^ are learnable parameter matrices. The coefficient α is a learnable residual coefficient. The hyperparameters *b* and *d* are set to *b* = 256 and *d* = 1536, respectively. The term 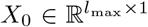represents the initial input cell pre-processed by the aforementioned workflow.

As performed in previous LLMs, we randomly mask the expressions of 20% of the genes (denoted as *M*) and then recover them based on the non-masked genes. Specifically, for the *M* masked gene expressions, we replace the gene expressions of the cell with a learnable weight vector X_*M*_ ∈ ℝ^1×*d*^ initialized to zero. Consequently, the feature *X*_*tmp*_ can be obtained as follows:

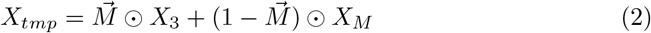

where ⊙ is the element-wise product, and 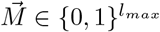 is the mask vector indicating the position of masked genes with value 0.

To learn the specific characteristic of each gene, we initialize a learnable embedding matrix *E*^*G*^ ∈ ℝ^24079×*d*^. The term 24079 represents the number of genes *ID* and the *d* = 1536 means the dimension of vector embeddings initialized based on each unique gene *ID*. We then integrate the gene expression and gene *ID* embeddings as follows:

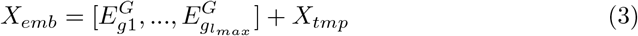

Furthermore, we incorporate an additional learnable weight *X*_*cls*_ ∈ ℝ^1×*d*^, which is appended to the gene expression embeddings. This weight facilitates the learning of cell-level features by aggregating gene information in the following manner:

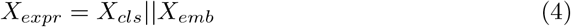

where the || symbol denotes the concatenation of two vectors.

### 4.4 The ERetNet module

CellFM learns gene embeddings and relationships from gene expression through the ERetNet module, a variant of the RetNet [34]. RetNet is an efficient, high-performance variant of the Transformer architecture with linear complexity. To better adapt largescale single-cell datasets, we have modified the RetNet module in three ways: First, we’ve increased the computational efficiency of the attention mechanism by replacing the Softmax function with the Relu activation function. Second, we’ve replaced the traditional feedforward network in RetNet with a gated bilinear network, which has led to improved model performance and a smoother training process. Third, we’ve refined the model’s training stability and performance by substituting the pre-layer LayerNorm in RetNet with the DeepNorm layer normalization technique [35]. Collectively, these modifications have resulted in the ERetNet module, which includes a Gated Multi-head Attention (MHA), a Gated Linear Unit (SGLU), and layer normalization (LN), all contributing to a more stable and effective gene expression analysis model.

#### 4.4.1 Gated Multi-head Attention (MHA)

The Gated Multi-head Attention (MHA) block is used to learn dependencies between genes, which is a variant of the Retention mechanism in the RetNet. To reduce the time-consuming exponential attention operations 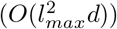 in the RetNet, we employ ReLU as the kernel function instead of the original Softmax computation of Q, K, and V in Retention to achieve a low time-consuming 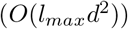. Furthermore, we scaled Q, K, and V following the study [36] to ensure the half-precision training as follows:

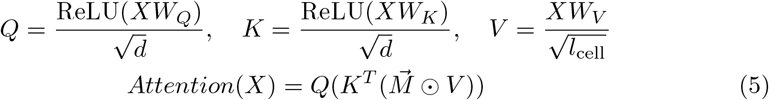

where *l*_*cell*_ = *l*_*max*_ + 1 denotes the number of expressed genes within each cell, 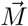 denotes the mask vector.

To enhance the CellFM’s representational power, we use *h* = *d*/*d*_*head*_ attention heads in each ERetNet layer, where *d*_*head*_ = 32 is the head dimension. Each head consists of three parameter matrices *W*_*Q*_, *W*_*K*_, 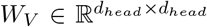. In addition, we add a swish gate [37] to increase the non-linearity of ERetNet layers. Formally, given input *X*_*expr*_, we define the MHA layer as follows:

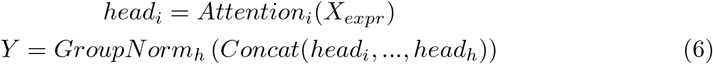

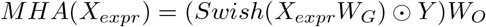

where *W*_*G*_, *W*_*O*_ ∈ ℝ^*d*×*d*^ are learnable parameters, and GroupNorm normalizes [38] the output of each head, following SubLN proposed in the study [39].

#### 4.4.2 Simple Gated Linear Unit (SGLU)

To improve model performance and a smoother training process, we’ve replaced the traditional feedforward network in RetNet with a gated bilinear network. The gated unit GLU introduces a multiplicative gating mechanism that explicitly indicates the model’s memory of each feature dimension, thereby smoothing the training process and facilitating better integration between channels. Considering that the gating mechanism inherently introduces nonlinear relationships, to further accelerate the model’s computation, this work, referring to literature [40], adopts the SGLU, which is based on the GLU formula [41] but omits the Swish activation function:

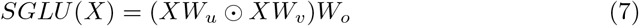

where ⊙ is the element-wise product.

#### 4.4.3 Layer normalization (LN)

The Transformer architecture in large single-cell models typically uses post-norm normalization after residual connections to enhance model depth and convergence. However, this can cause a gradient explosion as the model size grows. To counteract this, a pre-norm strategy is applied in RetNet for a stabilized training process, albeit with a potential performance trade-off.

To address this gap, CellFM employs the new post-norm normalization method DeepNorm [35]. DeepNorm reduces the contribution ratio of each network block to the output, thereby reducing the amount of gradient that needs to be updated and ensuring the stability of training.

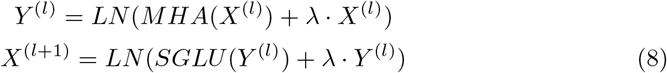

where LN(·) is LayerNorm and λ is a hyperparameter.

### 4.5 Low-Rank Adaptation (LoRA) module

Large models typically comprise hundreds of millions of parameters, resulting in considerable time consumption for full model training. To alleviate the burden of training on various datasets, we employ the Low-Rank Adaptation (LoRA) algorithm [42]. LoRA operates under the assumption that updates to pre-trained weights during fine-tuning can be low-rank decomposed. Hence, for a pre-trained weight matrix *W*_0_ ∈ ℝ^*n*×*k*^, we utilize low-rank decomposition to constrain the weight increment ∇*W* :

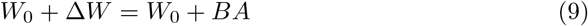

where *B* ∈ *R*^*d*×*r*^, *A* ∈ *R*^*r*×*k*^, and the rank *r* << *min*(*d, k*).

During the forward computation, both *W*_0_ and matrices *A* and *B* are used in calculations with the input *X*; however, during the backward propagation, the *W*_0_ parameter is frozen and does not undergo gradient updates, while only A and B are updated. It can be observed that in regular training, the training parameter count for *W*_0_ is *n* × *k*, whereas, with LoRA-based training, the training parameter count for *W*_0_ is the sum of the parameters of A and B, which is (*n* +*k*) × *r*. Since the dimension r is significantly smaller than *n* and *k*, the number of training parameters for the weights is greatly reduced, leading to a substantial decrease in computational overhead.

In the ERetNet architecture, the MHA has five weight matrices: *W*_*Q*_, *W*_*K*_, *W*_*V*_, *W*_*G*_, and *W*_*O*_, and the SGLU gating unit has three weight matrices: *W*_*u*_, *W*_*v*_, and *W*_*o*_. We consider the dimensions of these 8 matrices to be *d* × *d*. In this experiment, we limit the application of LoRA to only the ERetNet encoder part and freeze all model parameters for updates except for the weights of the LayerNorm layer.

### 4.6 Loss functions

#### Mean Squared Error (MSE)

CellFM is optimized to accurately predict the expression values of masked genes through the MSE loss. Specifically, we employ a fully connected MLP followed by the ERetNet module to estimate the expression value for *M* genes. The optimization of this objective involves utilizing the MSE loss at the masked positions, denoted as *M* mask. The MSE works as follows:

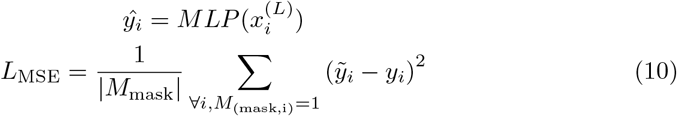

|*M*_mask_|denotes the count of ones in the mask gene vector *M*_*mask*_ for each cell. 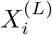 signifies the features derived from the ERetNet for gene *i* at the *L* layer.

To further enhance CellFM’s learning capabilities and channel aggregation, the feature corresponding to token *cls* is also leveraged, represented as 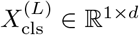. This feature passes through an additional network designed for predicting expression values. Specifically, a learnable parameter matrix *W*_cls_ ∈ ℝ^*d×d*^ and an activation function are employed. After mapping 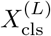, the result is multiplied with the embeddings of the genes to be predicted in the vocabulary *E*^*G*^ to compute inner products, yielding another set of predicted values. These are then compared with the actual values to calculate the mean squared error loss.

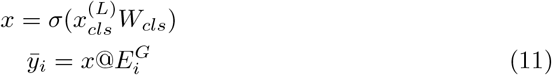

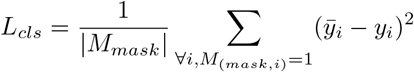

The @ symbol represents matrix multiplication. *σ* is an activation function Sigmoid. 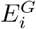 denotes the embedding of gene *i* in the vocabulary *E*^*G*^.

Finally, the total loss functions of CellFM can be obtained as follows:

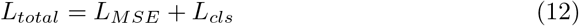

### 4.7 Implementation Details

CellFM consists of 40 stacked ERetNet blocks, with each block having 48 attention heads. During pre-training, we used approximately 100 million cells for training CellFM. The model was optimized using the Adam optimizer with a starting learning rate of 1e-7 and trained for a total of 2 epochs. The total batch size was 128, distributed equally across 4 Huawei Altas800 servers, each equipped with 8 Ascend910 NPUs. The MindSpore AI development framework powered the automatic data parallel training.

### 4.8 Pre-processing

The gene symbols across all raw count gene expression matrices were standardized using the reference mapping provided by the HUGO Gene Nomenclature Committee. This process included both human protein-coding genes and common mitochondrial genes, resulting in a comprehensive gene set G consisting of 24,078 genes. Ultimately, normalization and a log1p transformation were employed across all gene expression matrices to alleviate skewness in the data.

## Data availability

The evaluated datasets are accessible through the papers cited, with detailed link information available in the Supplementary Materials.

- Gene function prediction task: [[43]]
- Perturbation prediction task: [[44]]
- Cell annotation task: [[8]]
- Gene Network Analysis: [[44–46]].

## code availability

All codes used in this study can be available at: https://github.com/biomed-AI/CellFM.

## Funding and Acknowledgments

This study has been supported by the National Key R&D Program of China (2022YFF1203100), the National Natural Science Foundation of China (T2394502), the Research and Development Project of Pazhou Lab (Huangpu) [2023K0606], and the Postdoctoral Fellowship Program of CPSF (GZC20233321).

## Authors’ contributions

Y.Z., W.J., and Y.Y. conceived and supervised the project. Y.Z., Z.W., and C.X. contributed to the algorithm implementation. Y.Z., W.Y., and Y.Y. wrote the manuscript. Y.S., N.S., S.Y. C.Z., W.L., J.Z., N.F., H.Z., Y.L., J.F., and Y.Y. were involved in the discussion and proofreading.

